# Multimodal Characterization and Evolution of Malonate-induced Stroke Model: Advanced MRI, Histology-Molecular Profiling

**DOI:** 10.1101/2025.10.22.684053

**Authors:** Bayan El Amine, Benjamin Lemasson, Aurélien Delphin, Marc-Adrien Reveyaz, Nora Collomb, Emeline Lemarié, Sophie Bouyon, Antoine Boutin-Paradis, Hisham Altoufaily, Olivier Detante, Anne Briançon-Marjollet, Claire Rome

**Affiliations:** Univ. Grenoble Alpes, Inserm U1216, Grenoble Institut Neurosciences, 38000 Grenoble, France; Univ. Grenoble Alpes, Inserm U1300, CHU Grenoble Alpes, HP2 Laboratory, 38042 Grenoble cedex 9, France; Univ. Grenoble Alpes, Inserm, CNRS, CHU Grenoble Alpes, IRMaGe, 38000 Grenoble, France; Univ. Grenoble Alpes, CHU Grenoble Alpes, Neurology, Stroke Unit, 38043 Grenoble, France

**Author notes:** **Corresponding author** Dr Claire Rome, Grenoble Institut des Neurosciences (GIN) - U1216 Inserm UGA Eq 5 "Neuroimagerie Fonctionnelle et Perfusion Cérébrale" UGA - Site Santé, Chemin Fortuné Ferrini, 38700 La Tronche, France. Co-last authors (Anne Briançon-Marjollet, Claire Rome).

**Keywords:** Ischemic stroke, malonate, brain lesion, MRI

## Abstract

**Background:** Ischemic stroke is a leading cause of mortality and disability worldwide, yet therapeutic options remain limited. Preclinical models play a crucial role in understanding stroke pathophysiology and evaluating potential treatments. This study aimed to provide a comprehensive characterization of the temporal evolution of ischemic injury induced by malonate intracerebral injection using multiparametric magnetic resonance imaging (MRI) combined with histological and molecular analysis.

**Methods:** Focal ischemic lesions were induced by malonate injection in the striatum of rats. Lesion volume was monitored using T_2_-weighted MRI at multiple time points (Day 1, D7, D14, D28, and D56). Water content, reflecting vasogenic edema, was assessed *via* apparent diffusion coefficient (ADC) measurements, while vascular alterations were evaluated using blood volume fraction (BVF), vessel radius, and oxygen saturation (StO₂). Blood-brain barrier (BBB) permeability was quantified through gadolinium-enhanced MRI. Molecular analyses by RT-qPCR were conducted to assess oxidative stress, inflammation, and angiogenesis-related gene expression. Immunohistological staining was performed to investigate neuronal loss, astrocytic activation, and vascular remodeling.

**Results:** MRI analysis showed a significant and progressive decrease in lesion volume. Water content increased from D4 onward. Ischemic injury significantly altered vascular function, leading to increased vessel radius and BVF while reducing tissue oxygenation. BBB permeability was elevated at D7 and D56, accompanied by increased claudin-1 and aquaporin-1 expression. Molecular analysis revealed an upregulation of inflammatory markers (IL-6, TGF-β, NF-κB), oxidative stress response genes (SOD1, Nrf1), and impaired angiogenesis with increased Ang1/Ang2 but reduced VEGF/VEGFR1. Immunohistological analysis demonstrated neuronal loss, astrocytic activation, and vascular remodeling, characterized by increased ZO-1 and ColI-IV expression.

**Conclusion:** The observed changes in lesion volume, vascular function, inflammation, oxidative stress, and angiogenesis highlight key mechanisms underlying post-stroke recovery. These findings emphasize the importance of long-term monitoring in preclinical stroke models and may contribute to the development of novel therapeutic strategies.

## Introduction

Ischemic stroke is a major contributor to mortality, physical disability, and imposes a substantial socioeconomic burden worldwide. Its prevalence has continued to rise in recent years^1,2^. Ischemic strokes include subtypes like large artery atherosclerosis, cardioembolism, and cerebral small vessel disease (CSVD)^3^. Lacunar stroke, a key manifestation of CSVD, accounts for 20-30% of ischemic strokes and can present various syndromes based on lesion location. Silent lacunar strokes affect 20-50% of healthy elderly individuals^4^. These strokes are notable for a 20% recurrence rate, a 25% 5-year mortality rate, and associated morbidities like vascular cognitive impairment^5^.

Although extensive experimental and clinical research have been conducted over the past few decades, treatment options for ischemic stroke remain limited. Prompt restoration of cerebral blood flow using tissue plasminogen activator (tPA) and/or thrombectomy remains the most effective treatment. However, the therapeutic window for tPA is mainly restricted to first hours after stroke onset, meaning it is only applicable to approximately 5% of all patients. Complete recovery of neurological functions over the long term is rare^6^. Consequently, there is a critical need for alternative therapeutic strategies based on better pathophysiology knowledge.

Currently, no *in vitro* models exist that fully replicate the complex interactions between the vascular network, blood, and brain tissue during a stroke. Animal models of ischemic stroke provide unique opportunities to gain mechanistic insights and to assess the effects of therapeutic interventions on brain tissue *in vivo*.

In animal studies of ischemic stroke, infarct volume is considered the most precise metric, typically assessed post-mortem through histological methods. However, *in vivo* MRI offers the advantage of measuring not only the final infarct size but also tracking the temporal progression of the ischemic lesion. This longitudinal evaluation is a critical outcome measure, as functional recovery serves as the primary endpoint in clinical trials. One of the parameters monitored to assess this recovery can be the dynamic modification in microvessel function following a stroke, characterized through MRI follow-up, as demonstrated by Moisan *et al*.^7^.

Different animal models have been developed in the past several years to improve the understanding of brain ischemia, reflecting the diversity of brain injury outcomes following focal ischemia. Unlike large artery occlusive ischemic stroke, the unclear pathophysiological events in lacunar stroke make this subtype particularly challenging to model. However, the malonate-induced focal ischemia model can mimic some features of human lacunar stroke and is undoubtedly valuable for enhancing our understanding of the disease and testing treatments. Cirillo *et al.*^8^ established a rat model of focal ischemic injury through stereotaxic malonate injection. Malonate, a competitive inhibitor of the mitochondrial enzyme succinate dehydrogenase in the Krebs cycle, induces energy failure, excitotoxicity, and apoptotic cell death in both vascular and parenchymal tissues^9^. The malonate injection creates a lesion that results in measurable and reproducible sensorimotor deficits^8^. Furthermore, this preclinical model facilitates the generation of permanent, focal infarcts, with variability depending on topography. It is associated with low morbidity and high reproducibility, making it particularly useful for studying minor stroke (*ie* lacunar), the most common form of ischemic stroke^10^.

This study is the first to integrate longitudinal *in vivo* imaging with biological tools to characterize the temporal dynamics of biological response by MRI over a 56-day period following focal ischemic lesion at different time points, with a focus on molecular mechanisms during the transition phase, the subacute stage and the chronic phase^11^.

In rodents, stroke recovery is typically divided into distinct temporal phases based on underlying biological processes and functional changes. The acute phase (0–7 days post-stroke) is marked by infarct evolution, inflammation, edema, and cell death. This is followed by the subacute phase (approximately 7–28 days), characterized by heightened neuroplasticity, spontaneous recovery, and increased responsiveness to therapeutic interventions. The chronic phase begins beyond 28 days and is associated with stabilized deficits, reduced plasticity, and long-term remodeling processes. In the context of preclinical models, these time frames provide a framework for interpreting MRI-derived biomarkers and molecular dynamics across the transition from acute injury to long-term adaptation^12^.

The present study aimed to characterize the focal brain damage induced by malonate injection and assess the long-term consequences using multiparametric MRI combined with histological and molecular analysis, with a focus on characterizing the evolution of the ischemic lesion, angiogenesis and brain repair.

## Materials and Methods

### Animals

All animal procedures were run according to the French regulation with the approval of the local ethical committees and the French Ministry of Research (authorization #24116-2020012918159424 for GIN and #23039-2019112517102122 for HP2). Anesthesia was induced by inhalation of 5% isoflurane (Abbott Scandinavia AB, Sweden) in 20% O_2_ in air and maintained with 2-2.5% isoflurane through a facial mask (for surgical and imaging procedures). In all experiments, body temperature was monitored and maintained at 37.0 ± 0.5°C. Weight was measured regularly during the whole experiment (Figure S1).

### Malonate model and experimental groups

33 male Sprague Dawley rats (weight 280-300 g, 7 weeks old) were provided by Janvier labs (France). They underwent stereotactic surgery at D0 to induce brain ischemic injury by malonate injection. After local intradermal injection of 0.1 ml of 2% lidocaine and scalp incision, a 1 mm diameter hole was drilled in the skull at these stereotactic coordinates from bregma: AP -1.5 mm, ML +3.2 mm. Malonate (2 µl, 3M) was injected at a dorso-ventral depth of 5.3 mm^8^, targeting the striatum with a 5 μl-syringe (Hamilton 700 Series, Phymep, France, 250 μm gauge needle) connected to a micro-pump (injection rate: 0.33 μL/min). After the injection, the needle was left in place for an additional 4 min to avoid backflow. The burr hole was sealed with bone wax, skin resewn and rats were allowed to recover in their home cages. The rats were followed for 4 days (n=5), 7 days (n=5), 14 days (n=8), 28 days (n=8), and 56 days (n=7). To ensure the succession of the ischemic injury, a modified neurological severity score (mNSS) test adopted from ^13^ was performed on 4 rats before and 2 hours, 4 days, 7 days, 14 days, and 28 days after the surgery (Figure S2).

### *In vivo* MRI experiments

Overview of MRI timepoints and euthanasia timeline in the experimental rats is indicated in Table S1. MRI sessions (4.7T, Bruker Avance III, Germany, MRI facility of Grenoble IRMaGe, https://irmage.univ-grenoble-alpes.fr/) were set at D1, D4, D7, D14, D28 and D56 after malonate injection. Physiological parameters were monitored during MRI acquisition, including body temperature and respiratory rate. Breathing rate was kept constant (45-60 breaths/min) by modulating inhaled anesthesia. MRI sequences were performed as follows: T_2_ weighted (T_2_W) images (TR)/echo-time (TE) = 2200/36 ms, voxel size = 117×117×800 μm^3^, 19 slices, field of view (FOV) = 30×30 mm^2^ was performed to determine lesion volume. Diffusion-weighted, spin-echo-EPI (EPI-diff) (TR=2.2 s, TE=33 ms, 5 slices, FOV = 30×30 mm^2^, NA= 8, matrix = 128×128 and voxel size = 234×234×800 μm^3^) was acquired. This sequence was applied 6 times: three times without diffusion weighting and three times with diffusion weighting (b = 800 s.mm^-1^) in three orthogonal directions.

Blood volume fraction (BVF), vessel radius (R) and oxygen saturation (StO_2_) imaging were performed using a steady-state approach. Briefly, multiple gradient-echo sampling of the free induction decay and spin-echo (MGESFIDSE; TR = 4000 ms, 34 gradient-echoes between 2.3 and 92.3 ms, Spin-Echo = 60 ms; voxel size = 234×234×800 μm^3^, 5 slices, FOV = 30×30 mm^2^) were acquired before and after the injection of the UltraSmall SuperParamagnetic Iron Oxide nanoparticles (USPIO, P904®, Guerbet, France; 133 μmol Fe/kg body weight). A three-minute delay after the first injection was applied before starting the second MGEFIDSE acquisition.

Blood brain barrier (BBB) leakage was assessed using a dynamic contrast enhanced (DCE) sequence consisting of 25 of T_1_-weighted (T_1_W) spin-echo images (TR/TE = 800/4 ms, voxel size= 234×234×800 μm^3^, 5 slices, FOV= 30×30 mm^2^). The acquisition time of one repetition was 15.2 s. After the acquisition of four baseline images, a bolus of gadolinium Gd-DOTA (Dotarem®, Guerbet, France; 200 μmol/kg) was injected. Permeability-related parameters as well as time-to-peak and the percentage of signal enhancement were measured on the DCE signal curve during the first 3 min post-injection of the bolus.

### MRI Data Processing

The MRI data were reconstructed and analyzed using MP3 software^14^ developed in MATLAB (MathWorks, USA).

Apparent diffusion coefficient (ADC) maps were automatically computed on the Bruker scanner as the means of the ADCs observed in each of three orthogonal directions. BVF, R and StO_2_ were computed using the MR fingerprinting approach proposed by Delphin *et al.*^15^. Note that this dictionary was simulated for experiments using USPIO with a 200 μmol Fe/kg dose. BVF, R and StO_2_ reflect only the functional vessels. BBB permeability was calculated on the DCE images as the signal enhancement (SE, %) induced by Gd-DOTA extravasation.

For lesion volume, the region of interest (ROI) corresponding to the whole ischemic lesion was manually delineated on T_2_W images. The same volume was also delineated on the contralateral hemisphere as a control. Viable ROI was defined as a sub-region of the lesional ROI using these constraints: ADC = [0-2500 µm^2^.sec^-1^], R >0 µm; BVF >0 % and StO2 = [0-100 %]. The remaining non-viable part of the lesion is referred to as necrotic ROI. After extracting the brain using the PCNN3D algorithm ^16^, T_2_W anatomical images and lesion regions of interest were co-localised using the FLIRT function of FSL (the FMRIB Software Library^17^) for all animals imaged at D1 (n=33), and finally all co-localized lesions were added.

At the end of the MR acquisitions, anesthesia was induced by inhalation of 5% isoflurane (Abbott Scandinavia AB, Sweden) in 20% O_2_ in air through a facial mask and then animals were euthanized by decapitation through guillotine. Brains were collected, frozen in liquid nitrogen and stored at -80°C. Later, brains from D4, D7 and D28 animals were used for both molecular and immunohistological analysis.

### Gene Expression Analysis

Gene expression levels at D4, D7, and D28 were assessed using RT-qPCR as described (see *Supplementary Methods*).

### Immunohistology

Immunohistological staining was performed at D4, D7, and D28 to evaluate vascular, neuronal, and astrocytic markers (see *Supplementary Methods*).

### Dihydroethidium (DHE) and Cresyl Violet Staining

Oxidative stress and tissue integrity were assessed using DHE and cresyl violet staining, respectively (see *Supplementary Methods*).

### Statistical Analysis

For lesion volume, weight, mNSS, gene expression and immunohistological analysis, two-way ANOVA was used on GraphPad Prism software (Dotmatics, USA). For MRI results, we performed paired nonparametric Mann-Whitney tests to compare both hemispheres at each time point. Results are represented with a median with interquartile range. Any significant result was considered when p value <0.05. When a significant interaction was found between malonate and time effect in the two-way ANOVA, Fishers’ LSD post-hoc tests were performed and *p*-values at each time-point are reported in Table S4.

## Results

### General observations

Focal ischemia procedure was well tolerated by animals (33 rats). 7% of animals died during surgery because of technical problems, 3% during anesthesia for the follow-up MRI. All the surviving animals (90% of initial animals) were ambulatory, able to eat and drink independently, although they showed slight motor neurological deficits in the contralateral limbs.

### Lesion volume progressively decreases after ischemic injury

Cresyl violet staining was carried out to visualize the affected regions and assess the extent of the lesions following malonate-induced cerebral infarction. Malonate injection caused disruption to the cytoarchitecture of the striatum in all animals. The progression of the lesion volume was measured on T_2_W images obtained from MRI scans at D1, D7, D14, D28, and D56 following the induction of focal ischemia. The median lesion volume after malonate injection significantly decreased from 47 mm³ at D1 and 33 mm³ at D4, to 15 mm³ at D7 (*p*=0.0002), 12 mm³ at D14 (*p*<0.0001), 11 mm³ at D28 (*p*<0.0001), and 10 mm³ at D56 (*p*<0.0001), showing an important stabilization phase in the first 14 days after the lesion (Figure 1). The final mean infarct volume represented around 0.5% of the whole brain, and the cavity formed after the stroke occupied the core of the lesion. The selected coordinates used for malonate injection provided lesions to deep-brain structures. The lesion was successfully identified within the striatum, internal capsule, and thalamus. Notably, despite variations in lesion localization, a consistent and topographically coherent pattern of deep anatomical structure involvement was observed.

**Figure 1.**
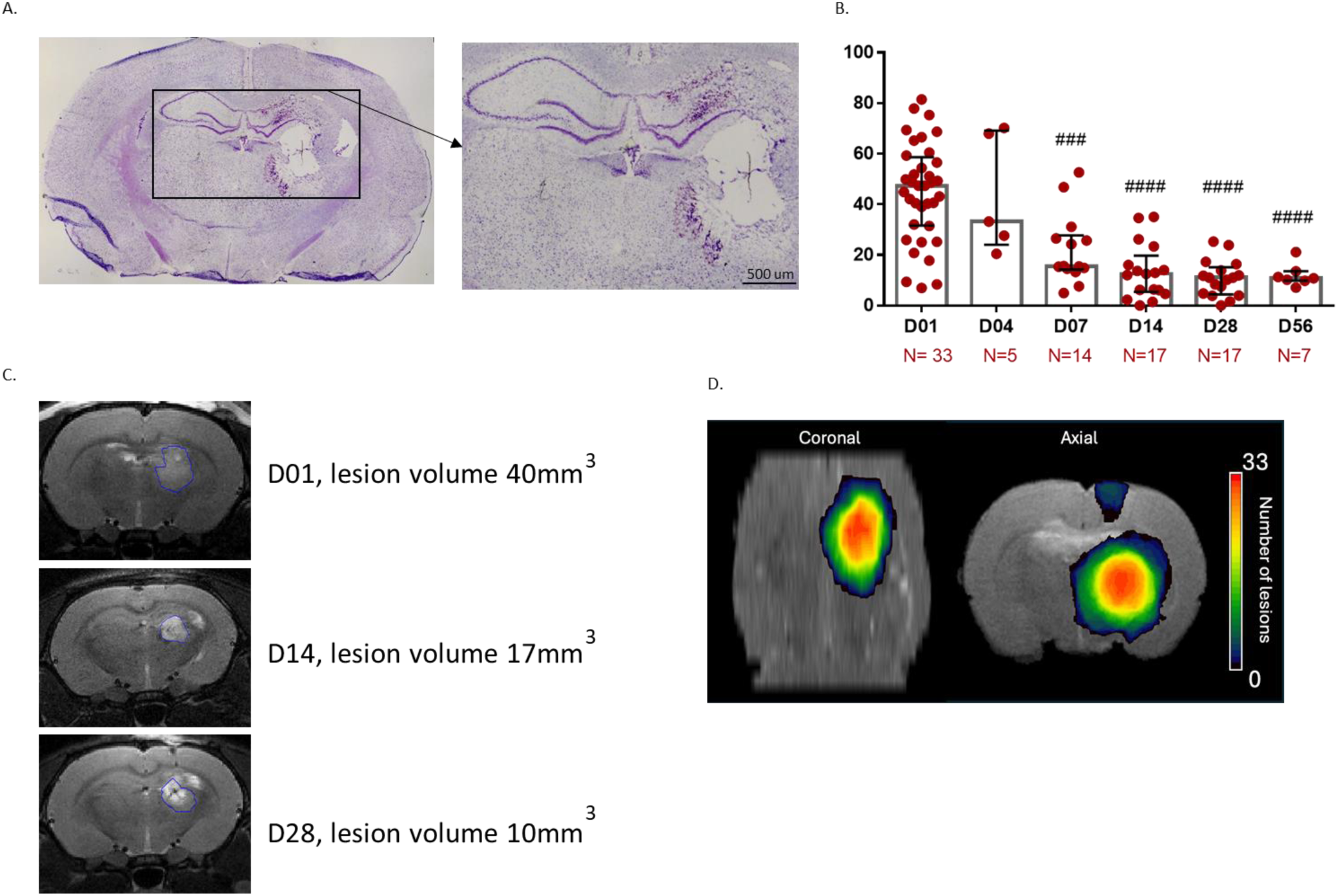
Brain lesion volume after 1, 4, 7, 14, 28 and 56 days post ischemic injury induced by malonate injection. A) Cresyl violet staining of the brain showing a developed lesion after ischemic injury at D1. B) The median lesion volume at different time points after ischemic injury showing that lesion size significantly decreased from its original size starting from D07 until D56. C) A representative figure for T_2_-weighted MRI images from one representative rat shows the lesion volume decreasing from 40 mm^3^ at D01 to 10 mm^3^ at D28. D) Overlap map with color-coded injured voxels from T_2_W anatomical images and lesion regions of interest at D1 (n=33), providing an overview of all the lesioned brain areas after injection of malonate. Data are expressed as median and interquartile range. ###:p=0.0001 and ####:p<0.0001 vs D1 (2-way ANOVA).

### Evaluation of water diffusion changes in ischemic injury

To assess the level of vasogenic edema within the viable ROI (Figure 2A), we used the ADC maps, which quantifies the magnitude of water molecule diffusion within the tissue (Figure 2B). At each time point, the contralateral hemispheres consistently showed an ADC of approximately 800 µm²/sec, representing the normal diffusion values of healthy tissue. In contrast, from D4 onward, the ADC in the ipsilateral hemispheres began to significantly increase, reflecting the development of vasogenic edema. Significant differences were observed between both hemispheres at all measured time points, except for D4.

**Figure 2.**
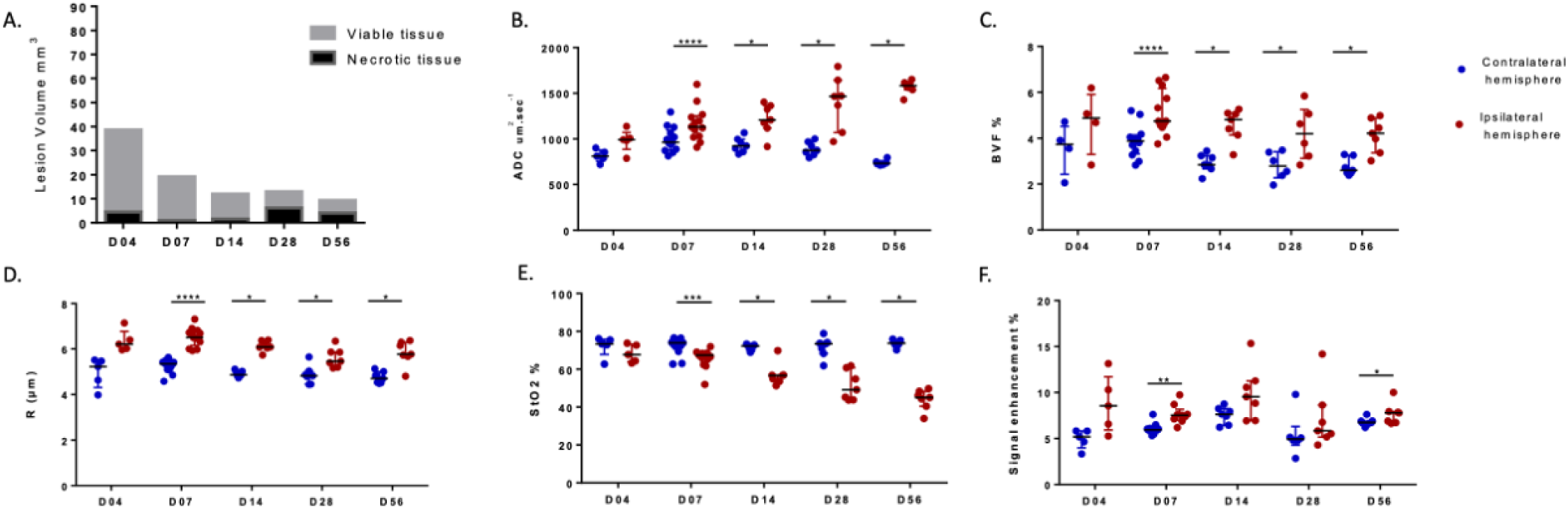
Multiparametric MRI. A. The percentage of viable and necrotic tissue with respect to the total lesion volume. B.The evolution of vessel radius (R). C. Blood volume fraction (BVF). D. Oxygen saturation (StO_2_) E. Signal enhancement F. Apparent diffusion coefficient (ADC) observed by MRI at D4, D7, D14, D28, and D56 after ischemic injury by malonate injection. n=5 at D4, n=13 at D7, n=8 at D14, n=8 at D28 and n=7 at D56. Data is represented by medians and IQR. *: 0.03>p>0.01, **: 0.005>p>0.001, ***: 0.0005>p>0.0001, ****: p<0.0001 vs contralateral hemisphere, paired non-parametric Mann-Whitney test.

### Blood volume, vessel radius, and tissue oxygen saturation are altered after ischemic injury

We measured BVF(%), R (µm), and StO_2_(%) at 4, 7, 14, 28, and 56 days following malonate injection. Compared to the contralateral hemisphere, ischemic injury induced an increase of R and BVF, along with a decreased tissue oxygenation in the viable ROI of the ipsilateral hemisphere in all time points except for D4 (Figure 2C, D, E). No significant changes were observed in the value of R, BVF, StO_2_ and ADC in the contralateral hemisphere over the 56-days period (Figure 2B-E).

### Blood brain barrier permeability is increased by ischemic injury

Compared to the contralateral hemisphere, BBB permeability was significantly increased in the ischemic lesion at D7 and D56, with no notable changes at other time points and in the contralateral hemisphere (Figure 2F). Moreover, molecular analysis showed that after ischemic injury, the gene expression of the tight junction protein claudin 1 and the water channel aquaporin 1 increased at D7 compared to contralateral hemisphere (Figure 3A), without significant changes in other tight junctions (claudin 5 and occludin; Figure 3A) or BBB transporters (P-gp and MRP1; Figure 3B). Additionally, the ipsilateral hemisphere exhibited significant changes between consecutive days in claudin 1, aquaporin 1, P-gp and MRP1 indicating a time-dependent effect of the ischemic injury (See Table S4).

**Figure 3.**
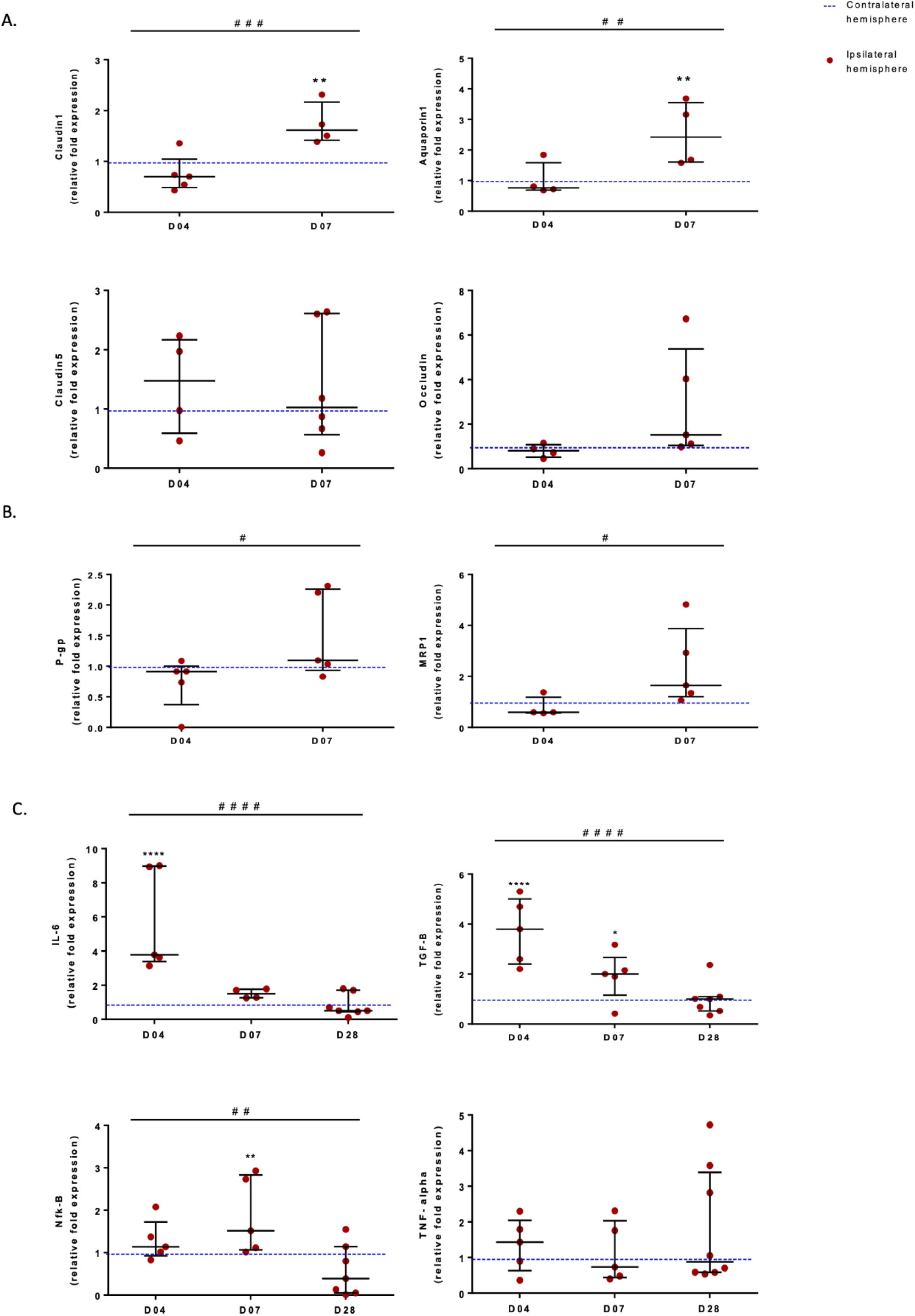
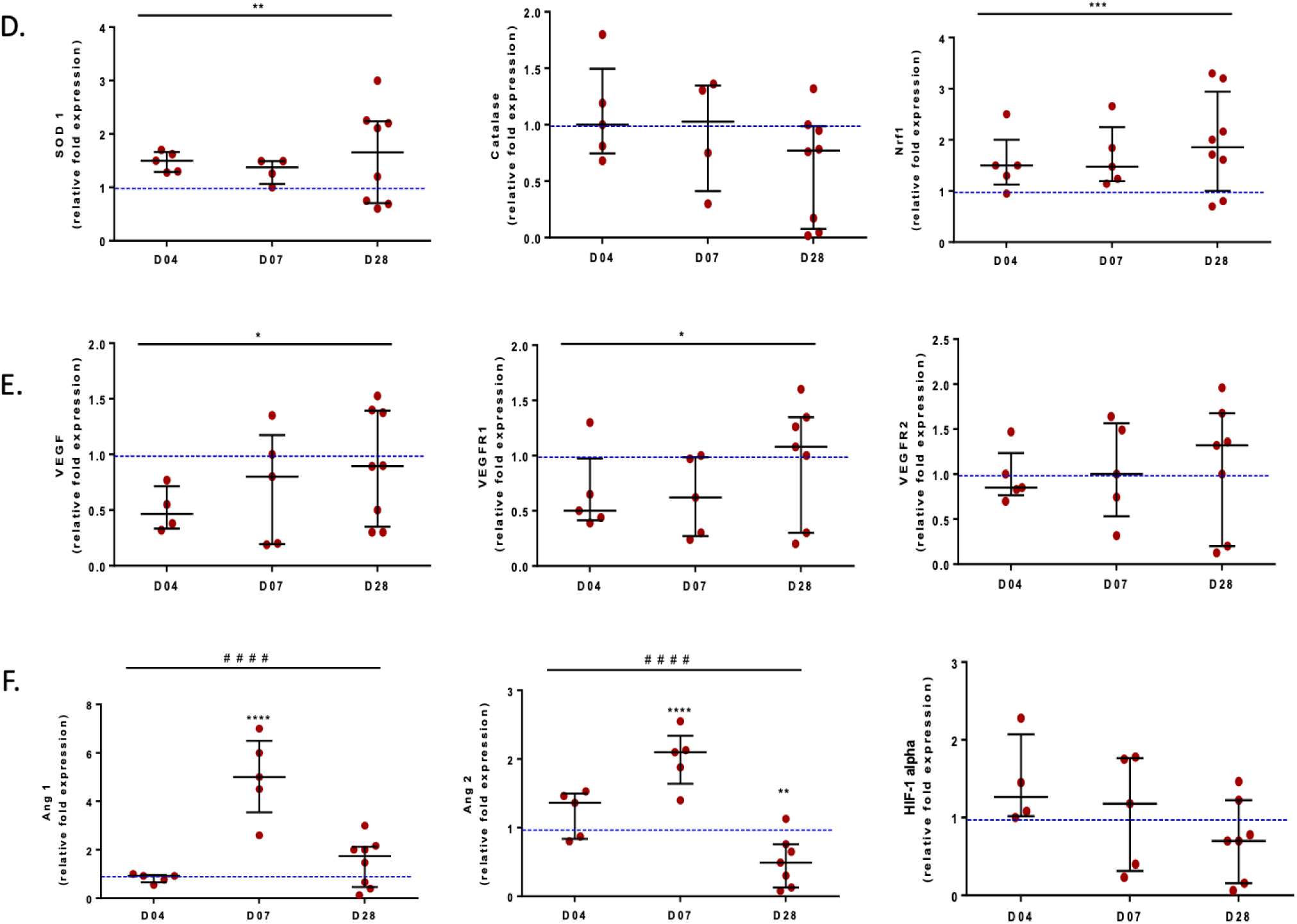
Brain mRNA expression of 19 genes after ischemic injury. The quantitative values are expressed as ratios of values in the lesioned hemisphere normalized to the contralateral normal tissue (dotted line). n=5 at D4, n=5 at D7, and n=7-8 for D28. Data are represented by medians and IQR. */#: 0.03>p>0.01, **/##: 0.005>p>0.001, ***/###: 0.0005>p>0.0001, ****/####: p<0.0001, 2-way ANOVA. * indicates the difference between ipsilateral and contralateral hemisphere, # indicates the difference between days.

### Ischemic injury increased brain inflammation, oxidative stress and impaired angiogenesis

The expression levels of various genes were monitored over time (D4, D7, and D28) at the site of the ischemic lesion using reverse transcription-quantitative PCR (RT-qPCR). These genes were associated with inflammation (IL-6, TGF-β, TNF-α, NF-κB, Figure 3C), oxidative stress (SOD1, catalase, and Nrf1, Figure 3D), growth factors and angiogenesis (VEGF, VEGFR1, VEGFR2, Ang1, and Ang2, Figure 3E and F), and hypoxia (HIF-1ɑ, Figure 3F).

Ischemic injury led to a significant increase in inflammation in ipsilateral hemisphere, as evidenced by elevated level of IL-6 (p=0.0004), TGF-β (p<0.0001) at D4 and NF-kB (p=0.08) at D7 without significant effect on TNF-𝛼. Additionally, independent of the time, the cellular adaptive response to oxidative stress was enhanced with increase in both SOD1 (p=0.006) and Nrf1 (p=0.007) while catalase (Figure 3D) or DHE levels remained unaffected (Figure 4A). Ischemic injury also impaired angiogenesis by increasing Ang1 (p<0.0001) and Ang2 (p<0.0001) at D7 and decreased both VEGF (p=0.0121) and VEGFR1 (p=0.0257) without significant effect on VEGFR2 (Figure 3 E-F). Finally, during the entire follow-up, no significant difference in the levels of HIF-1ɑ was observed between the contralateral and ipsilateral hemispheres (Figure 3F).

**Figure 4.**
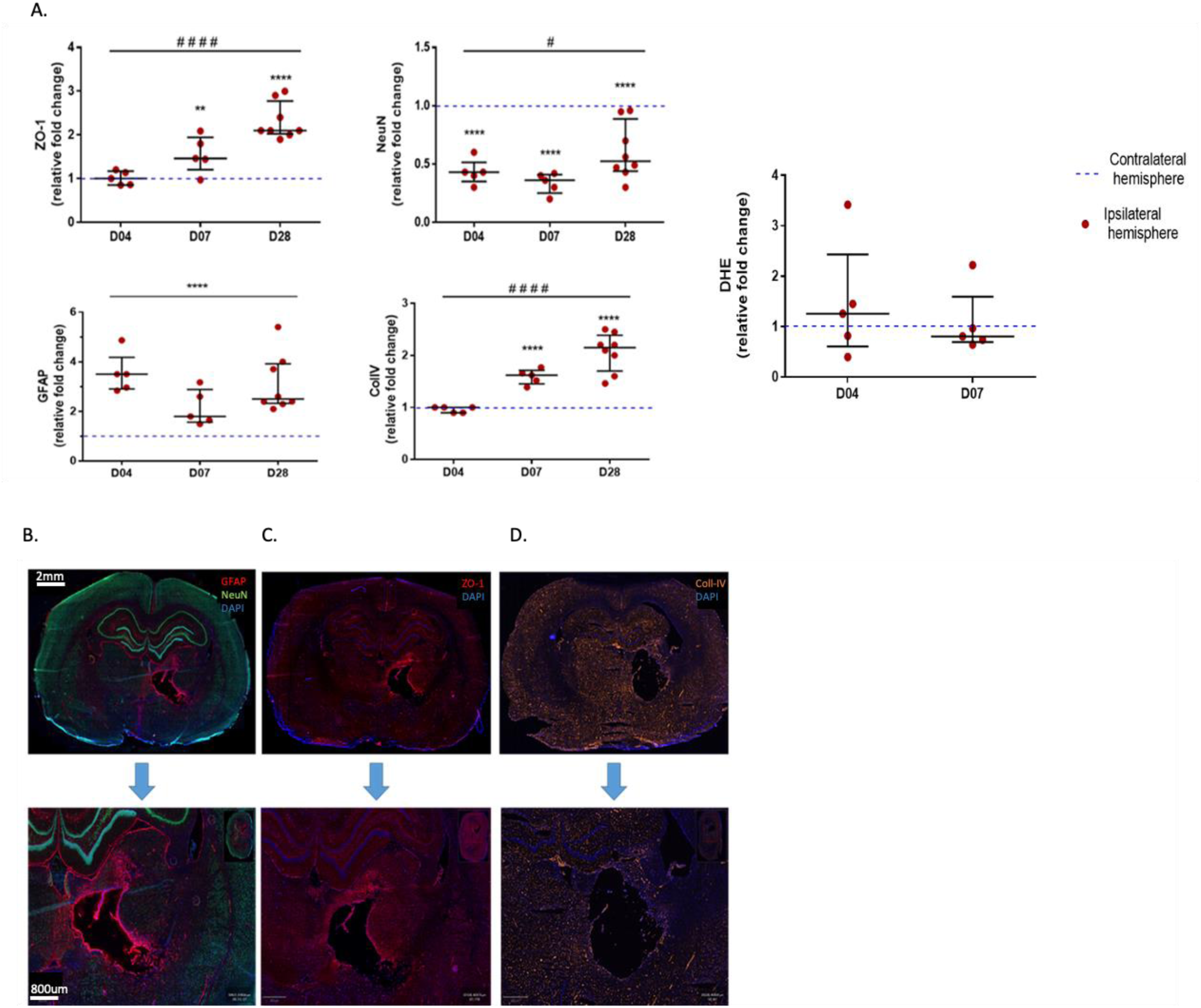
Immunohistological analysis of brains after ischemic stroke. A. Relative expression of NeuN, ZO-1, GFAP, ColI-IV in the viable part of the lesion and DHE in the perilesional area. B. NeuN (green), GFAP (red) and DAPI (4′,6-diamidino-2-phenylindole, Blue). C. ZO-1 (red) and DAPI (blue). D. ColI-IV (yellow) and DAPI (blue). n=5 at D4, n=5 at D7, and n=8 for D28. Data is represented by median with interquartile range. */#: 0.03>p>0.01, ***/###: 0.0005>p>0.0001, ****/####: p<0.0001 for ipsilateral vs contralateral hemisphere, 2-way ANOVA. * indicates the difference between ipsilateral and contralateral hemisphere, # indicates the difference between days.

Moreover, the ipsilateral hemisphere exhibited significant changes between consecutive days in some markers (IL-6, TGF-B, NF-𝜅B, Ang1 and Ang2), indicating a time-dependent effect of the ischemic injury (See Table S4).

### Ischemic injury induces brain neuronal loss, astrocytic activation and vascular remodeling

To characterize neuronal loss (NeuN), astrocytic activation (GFAP) and brain vasculature (ColI-IV and ZO-1), immunohistological analysis were conducted on brain sections at 4, 7, and 28 days post-ischemic injury (Figure 4).

The results revealed a decrease in the number of neurons (NeuN+ cells) (p<0.0001) and an increase in astrocyte activation (GFAP+ cells) (p<0.0001) in the ipsilateral hemisphere. Furthermore, ischemic injury also triggered vascular remodeling, as evidenced by an increase in both tight junction ZO-1 (p<0.0001) and the extracellular matrix component ColI-IV (p<0.0001) in the ipsilateral hemisphere. Additionally, the ipsilateral hemisphere exhibited significant increases between consecutive days in all the markers, except for GFAP, indicating a time-dependent effect of the ischemic injury (See Table S4).

Additionally, DHE staining was performed at D4 and D7 to monitor superoxide production. Four days after the induction of the brain lesion, no significant difference of staining was observed in the perilesional area when compared to the contralateral hemisphere.

## Discussion

The clinical heterogeneity of stroke complicates research, necessitating large patient cohorts to minimize confounding factors. Experimental focal cerebral ischemia models, particularly in rodents, offer a controlled environment that allows researchers to investigate specific pathogenic events following ischemic stroke and explore novel therapeutic strategies. In this study, we characterized the focal minor stroke developed by Cirillo *et al.*^8^. In this model, the stereotaxic injection of the mitochondrial toxin malonate induces focal deep damage in the striatum area. A key advantage of the malonate model, in comparison to other existing models, lies in its minimal invasiveness, which avoids the induction of artificial surgery-induced systemic inflammation. Additionally, it offers low variability in lesion location through precise stereotactic malonate administration and is effective in inducing a focal ischemic lesion confined to the targeted structures. Consistent with the literature on the malonate model in rats, this model is characterized by considerate reproducibility and favorable survival outcomes with ∼5% mortality rate. To our knowledge, this study is the first to integrate longitudinal *in vivo* imaging with biological tools to characterize the temporal dynamics of biological response by MRI over a 56-day period following focal cerebral and ischemic lesion at different time points, with a focus on molecular mechanisms during the transition phase, the subacute stage and the chronic phase^11^. We associated molecular data with MRI analysis to characterize the lesion evolution.

### The Malonate Model as a Model of Lacunar Stroke

Our studies confirmed that malonate injection caused progressive structural changes in the lesion core in the following days of observation. We observed that the lesion size was largest before D4. Indeed, the lesion volume, monitored by MRI, showed a decrease from day 1 to day 28 and a stabilization during the weeks after. This lesion location mimics stroke formed by occlusion of the deep perforating arteries. The work described here focused on injury to the deep-brain structures, since a relatively small lesion causes clinical symptoms^19^, considering the mNSS results, thus categorizing this model as a minor stroke model^20^ as previously defined in other studies^8,21^. Furthermore, we have confirmed that this model is not only applicable and reliable but also rapid and highly effective in inducing focal ischemic-type lesions confined to deep brain structures. This model can therefore reduce the number of animals used while maintaining statistical significance in therapeutic studies. It is important to note that to ensure meaningful comparisons of therapeutic effects, it is crucial to have comparable pre-treatment groups.

### Characterization of the Model During the Transition, Subacute and Chronic Phases

The evolution of MRI parameters reflecting the pathophysiology of stroke has been studied to some extent in humans and in animal models. Various MRI parameters that reflect the pathology of ischemic brain tissue differently evolve and can be used to estimate the time elapsed after stroke onset^7^. The malonate model is described as being suitable for studies on subacute and long-term lesions, but less so for hyperacute and acute phase studies, as it does not replicate vascular occlusion/reperfusion better modeled by thrombo-embolic models^8^. However, after the hyperacute post-ischemic phase, a "transition to the subacute phase," occurring between days 3 and 7, has been well-documented. This phase is characterized by a complex molecular expression pattern that is crucial for post-stroke recovery^7^. The MRI data, in conjunction with molecular data, enable a comprehensive characterization of the malonate model during the latter part of the acute/subacute phase and the chronic phase. We firstly characterized the processes occurring during the transition phase in this model.

During the transition phase (D3 to D7), we observe an increase in vascular permeability, which can be correlated with the upregulation of inflammatory factors such as IL-6, NF-κB, and TGF-β (mediators of post-ischemic inflammation)^22^ and oxidative stress (SOD1 and NRF1 at D4). By contrast, this increase of permeability is not associated with VEGF because we do not observe any overexpression of VEGF, which is underexpressed throughout the entire observation period. During this period, we also observed a reduction in vascular density, due to an increase in both the BVF and cerebral vessels radius, alongside the increase in BBB permeability. These findings are in alignment with similar observations made in three-vessel occlusion^23^ and MCAo models^7^, underscoring the consistency of vascular alterations across different ischemic models. Following ischemic stroke, BBB disruption, coupled with the altered interaction between vascular endothelial cells and the vascular basement membrane, facilitates the infiltration of neurotoxic substances into the brain parenchyma, leading to irreversible tissue damage^24^. However, it is important to note that the increase in permeability observed in our model is less pronounced than in the other models. The increase in permeability observed during the transition phase is supported by various molecular markers, including the overexpression of IL-6, Claudin-1 and AQP1 in the ipsilateral hemisphere. These findings are consistent with previous stroke research, which reports a rapid elevation of inflammatory cytokines, such as IL-6, following ischemic injury^25^. This elevation results in exacerbated BBB breakdown. Additionally, the overexpression of Claudin-1 has been associated with the disruption of the BBB after ischemic stroke, a process linked to the destabilization of tight junction (TJ) complexes. After the acute phase of stroke, Claudin-1 is expressed in the most permeable vessels, where it integrates into the TJs through direct interaction with ZO-1, which is also overexpressed under these conditions in the ipsilateral hemisphere^26^. Moreover, the BBB disruption diminishes during the subacute stage, correlating with an increase in ZO-1 expression on D7. Furthermore, molecular analyses revealed a pattern that is consistent with previous research^7,11^.

P-glycoprotein (P-gp) and MRP1 are two ATP-dependent transmembrane glycoproteins. P-gp is primarily expressed on the luminal surface of brain microvascular endothelial cells that form the BBB and MRP1 is predominantly situated at the basolateral membrane of the choroidal epithelium, facing the stromal/blood space^27^. Increased P-gp expression has been observed in capillary endothelial cells following focal cerebral ischemia^28^, and elevated P-gp levels after ischemic stroke^29^ have been shown to worsen the breakdown of the BBB rapidly and transiently within 24 hours post-injury. Concerning MRP1, it has been reported that its expression decreases in response to focal cerebral ischemia during the 72 hours following the injury but was still functional in the brain after ischemia^30^. However, it is worth noting that it has been demonstrated that the deactivation of P-gp and MRP1 does not affect post-ischemic neurological deficits, secondary neurodegeneration, or neurogenesis^31^. In our study, focusing on the transition and subacute phases, we observed that the expression level of P-gp and MRP1 remained stable in the ipsilateral hemisphere (96h post-ischemia induction) compared to contralateral hemisphere, consistent with findings from previous studies.

Moreover, an increase in AQP1 mRNA expression was observed during this period. Interestingly, AQP1, though highly expressed in peripheral endothelial cells, is typically absent in the endothelium of normal cerebral capillaries^32^. However, AQP1 has been shown to be significantly expressed in choroid plexus endothelial cells during vasogenic edema following cerebral ischemia^33^. In stroke patients, AQP1 has been identified in both perivascular astrocytes and the brain parenchyma, where it localizes in the glial scar and the peri-liquefaction zone surrounding the lesion^34^. Additionally, AQP1 is implicated in critical processes such as angiogenesis^35^. This is of particular interest, as angiogenesis is known to occur within 4 to 7 days following cerebral ischemia at the edge of the ischemic core^36^. This process underscores the potential role of AQP1 in the repair and remodeling of the brain after ischemic injury.

In this context, focusing on the remodeling of the functional microvascular system is crucial for advancing stroke treatment^7,37^. VEGF acts as a key molecular hub in vascular remodeling, while angiopoietin (Ang), which is upregulated in the later stages of remodeling, promotes vascular endothelial cell maturation and creates a stable environment for angiogenesis^38^. Furthermore, VEGF and Ang have been shown to work in concert to facilitate vascular remodeling in the ischemic penumbra during stroke^39^. On day 4, we observed a drop in the expression of VEGF and VEGFR, in line with previous reports^40^ alongside a significant increase in TGF-β expression, which then declined by day 28. This molecular angiogenic pattern signifies the ‘transition stage’ between the acute (D1–D3) and subacute (D7–D25) stages of recovery^11^. The ‘Transition stage’ (D3-D7) involves TGF-β and Ang1, which are in correlation with ColI-IV evolution.

Notably, the increased levels of TGF-β observed in the infarcted brain may also indicate the initiation of an anti-inflammatory signaling pathway during the early stages of stroke. It has been reported that TGF-β is upregulated following ischemia, primarily in microglia and macrophages and TGF-β signaling represents a distinct mechanism by which astrocytes mitigate neuroinflammation, with TGF-β being an anti-inflammatory and neuroprotective cytokine that is upregulated in the subacute phase post-stroke^41^. Beyond its neuroprotective effects, TGF-β modulates immune cells by suppressing inflammation through the inhibition of Th1 and Th2 responses and by promoting the development of Treg cells^42^. Thus, post-ischemic TGF-β production aids tissue repair by resolving inflammation and providing direct cytoprotection to surviving cells in the ischemic region.

TNF-α/NF-𝜅B-mediated neuroinflammatory responses within the brain parenchyma are critical in the disruption of the BBB, primarily through the induction of vasogenic edema via NF-𝜅B activation. In this study, since no reperfusion occurs, vasogenic edema is present but we observed an absence of changes in TNF-α expression at times studied^41^. These results are consistent with those presented in Nishi *et al.*^43^, where an increase in TNF-α was observed only within the first 24 hours after focal ischemia.

In the context of ischemia, measurements of the ADC and StO_2_ revealed increasing tissue necrosis within the ipsilateral hemisphere, further substantiating the progressive nature of ischemic injury, to D28 and a stabilization of the lesion to D56. A rapid and stable reduction in NeuN expression, indicative of neuronal damage or loss, was observed, which was concomitant with an upregulation of GFAP expression, signaling reactive astrocytosis and NF-𝜅B, which contributes to neuronal cell death in cerebral ischemia^44^. NeuN, a marker for mature neurons, serves as an indicator of neurodegeneration or impaired neuronal survival, both of which are frequent outcomes of ischemic insults. These findings align with results from other ischemic models, thereby reinforcing the role of astrocytes in neuroinflammation and tissue repair^45^.

### Differences with reperfusion models

In cerebral ischemia and reperfusion models, cerebral blood flow is impaired due to vessel occlusion in regions of the brain that are normally oxygenated. Reoxygenation during reperfusion triggers various enzymatic oxidation reactions. Under normal physiological conditions, mitochondria generate superoxide anion radicals and hydrogen peroxide^46^. H_2_O_2_ formation through Fenton’s reaction is known to induce BBB leakage^47^. In numerous studies, it has been established that oxidative stress is a critical factor in mediating cell death following cerebral ischemia-reperfusion (I/R), during which the elevated production of reactive oxygen species (ROS) leads to substantial cerebral tissue damage^48^. Moreover, antioxidant enzymes, including SOD and catalase (CAT), are crucial in defending against oxidative stress following acute ischemia. These enzymes are the primary means by which cells counteract ROS. Cytoplasmic SOD1 is known to have protective effects against I/R injury. Post-ischemia, dysfunctional mitochondria generate excess ROS, initiating a vicious cycle where ROS-induced damage further promotes ROS production. Both SOD and CAT scavenge ROS, providing neuroprotection from ischemic damage^49^. SOD1, as a critical indicator of antioxidative stress defense, reduces neuronal loss and limits the extent of injury during the acute post-ischemic period^48^. In our model and within the established timeframes, it is important to note that the malonate-induced lesion induced a moderate oxidative stress. Indeed, adaptative antioxidant activity is observed, as evidenced by a stable increase in the expression of SOD1 and Nrf1 over time but without affecting catalase expression or DHE staining. This limited oxidative response could be associated with the absence of reperfusion, which is responsible for the formation of H_2_O_2_.

## Conclusion

This study investigates the focal lacunar stroke model induced by minimally invasive intracerebral injection of malonate, a mitochondrial toxin, offering a method to explore ischemic stroke. The model induces controlled, reproducible focal damage in deep brain structures, simulating stroke caused by perforating artery occlusion. The results highlight significant molecular changes, including increased vascular permeability, upregulation of inflammatory pathways (IL-6, NF-kB, TGF-β), and altered expression of TJ proteins such as Claudin-1, indicating BBB disruption. Notably, TGF-β signaling plays a key role in neuroprotection and tissue repair. The study also observes that molecular changes underlying vascular remodeling and neuroinflammation correlate with lesion progression. Compared to reperfusion models, this model shows lower injury severity, limited oxidative stress, but higher reproducibility and lower mortality which makes it a more raffinated model in terms of ethics. This underscores the relevance of the malonate model as a valuable tool for investigating the molecular mechanisms underlying ischemic injury and recovery.

## Non-standard Abbreviations and Acronyms

ADC: apparent diffusion coefficient
Ang1: angiopoietin 1
Ang2: angiopoietin 2
BBB: blood-brain barrier
BVF: blood volume fraction
ColI-IV: collagen type IV
DAPI: 4′,6-diamidino-2-phenylindole
DHE: dihydroethidium
FOV: field of view
FSL: FMRIB Software Library
GFAP: glial fibrillary acidic protein
HIF-1: hypoxia inducible factor 1
IL-6: interleukin 6
IV: intravenous injection
MRI: magnetic resonance imaging
MRP1: multidrug resistance-associated protein 1
NeuN: neuronal nuclear antigen
Nf-kB: nuclear factor kappa light chain enhancer of activated B cells
Nrf1: nuclear respiratory factor 1
PBS: phosphate-buffered saline
P-GP: p-glycoprotein
R: vessels radius
ROI: region of interest
RT-qPCR: reverse transcription-quantitative polymerase chain reaction
SE: signal enhancement
SOD1: superoxide dismutase 1
StO_2_: oxygen saturation
T_1_W: T_1_ weighted
T_2_W: T_2_ weighted
TGF-B: transforming growth factor beta
tMCAO: transient middle cerebral artery occlusion
TNF-alpha: tumor necrosis factor alpha
VEGF: vascular endothelial growth factor
VEGFR1: vascular endothelial growth factor receptor 1
VEGFR2: vascular endothelial growth factor receptor 2
ZO-1: Zona occludens 1

## Acknowledgments

The authors thank both Nacera Bakdouri for her assistance during the *in vivo* experiments, and the animal facility working team for their assistance in animal care and housing at the GIN and PHTA animal facilities. This work was supported by the Photonic Imaging Center of Grenoble Institute Neuroscience (Univ Grenoble Alpes– Inserm U1216) which is part of the ISdV core facility and certified by the IBiSA label. This work was performed on the IRMaGe platform member of France Life Imaging network (grant ANR-11-INBS-0006).

## References

1. GBD 2019 Stroke Collaborators. Global, regional, and national burden of stroke and its risk factors, 1990–2019: a systematic analysis for the Global Burden of Disease Study 2019. Lancet Neurol. 2021;20:795–820.

2. Tsao CW, Aday AW, Almarzooq ZI, Anderson CAM, Arora P, Avery CL, Baker-Smith CM, Beaton AZ, Boehme AK, Buxton AE, et al. Heart Disease and Stroke Statistics-2023 Update: A Report From the American Heart Association. Circulation. 2023;147:e93–e621.

3. Caplan LR. Lacunar Infarction and Small Vessel Disease: Pathology and Pathophysiology. J Stroke. 2015;17:2–6.

4. Bamford J, Sandercock P, Jones L, Warlow C. The natural history of lacunar infarction: the Oxfordshire Community Stroke Project. Stroke. 1987;18:545–551.

5. Vermeer SE, Longstreth WT, Koudstaal PJ. Silent brain infarcts: a systematic review. Lancet Neurol. 2007;6:611–619.

6. Ahmed N, Wahlgren N, Grond M, Hennerici M, Lees KR, Mikulik R, Parsons M, Roine RO, Toni D, Ringleb P, et al. Implementation and outcome of thrombolysis with alteplase 3-4.5 h after an acute stroke: an updated analysis from SITS-ISTR. Lancet Neurol. 2010;9:866–874.

7. Moisan A, Favre IM, Rome C, Grillon E, Naegele B, Barbieux M, De Fraipont F, Richard M-J, Barbier EL, Rémy C, et al. Microvascular Plasticity After Experimental Stroke: A Molecular and MRI Study. Cerebrovascular Diseases. 2014;38:344–353.

8. Cirillo C, Le Friec A, Frisach I, Darmana R, Robert L, Desmoulin F, Loubinoux I. Focal Malonate Injection Into the Internal Capsule of Rats as a Model of Lacunar Stroke. Front Neurol. 2018;9:1072.

9. Vaysse L, Conchou F, Demain B, Davoust C, Plas B, Ruggieri C, Benkaddour M, Simonetta-Moreau M, Loubinoux I. Strength and fine dexterity recovery profiles after a primary motor cortex insult and effect of a neuronal cell graft. Behav Neurosci. 2015;129:423–434.

10. Maïer B, Kubis N. Hypertension and Its Impact on Stroke Recovery: From a Vascular to a Parenchymal Overview. Neural Plast. 2019;2019:6843895.

11. Arai K, Jin G, Navaratna D, Lo EH. Brain angiogenesis in developmental and pathological processes: neurovascular injury and angiogenic recovery after stroke. FEBS J. 2009;276:4644–4652.

12. Cirillo C, Brihmat N, Castel-Lacanal E, Le Friec A, Barbieux-Guillot M, Raposo N, Pariente J, Viguier A, Simonetta-Moreau M, Albucher J-F, et al. Post-stroke remodeling processes in animal models and humans. J Cereb Blood Flow Metab. 2020;40:3–22.

13. Chen J, Sanberg PR, Li Y, Wang L, Lu M, Willing AE, Sanchez-Ramos J, Chopp M. Intravenous administration of human umbilical cord blood reduces behavioral deficits after stroke in rats. Stroke. 2001;32:2682–2688.

14. Brossard C, Montigon O, Boux F, Delphin A, Christen T, Barbier EL, Lemasson B. MP3: Medical Software for Processing Multi-Parametric Images Pipelines. Front Neuroinform. 2020;14:594799.

15. Delphin A, Boux F, Brossard C, Coudert T, Warnking JM, Lemasson B, Barbier EL, Christen T. Enhancing MR vascular Fingerprinting with realistic microvascular geometries. Imaging Neuroscience. 2024;2:1–13.

16. Chou N, Wu J, Bai Bingren J, Qiu A, Chuang K-H. Robust automatic rodent brain extraction using 3-D pulse-coupled neural networks (PCNN). IEEE Trans Image Process. 2011;20:2554–2564.

17. Jenkinson M, Beckmann CF, Behrens TEJ, Woolrich MW, Smith SM. FSL. NeuroImage. 2012;62:782–790.

18. Gaucher J, Montellier E, Vial G, Chuffart F, Guellerin M, Bouyon S, Lemarie E, Yamaryo-Botté Y, Dirani A, Ben Messaoud R, et al. Long-term intermittent hypoxia in mice induces inflammatory pathways implicated in sleep apnea and steatohepatitis in humans. iScience. 2024;27:108837.

19. Saa JP, Tse T, Koh GC-H, Yap P, Baum CM, Uribe-Rivera DE, Windecker SM, Ma H, Davis SM, Donnan GA, et al. Characterization and individual-level prediction of cognitive state in the first year after ‘mild’ stroke. PLOS ONE. 2024;19:e0308103.

20. Yaghi S, Herber C, Boehme AK, Andrews H, Willey JZ, Rostanski SK, Siket M, Jayaraman MV, McTaggart RA, Furie KL, et al. The Association between Diffusion MRI-Defined Infarct Volume and NIHSS Score in Patients with Minor Acute Stroke. J Neuroimaging. 2017;27:388–391.

21. Colitti N, Desmoulin F, Le Friec A, Labriji W, Robert L, Michaux A, Conchou F, Cirillo C, Loubinoux I. Long-Term Intranasal Nerve Growth Factor Treatment Favors Neuron Formation in de novo Brain Tissue. Front Cell Neurosci. 2022;16:871532.

22. Chamorro Á, Meisel A, Planas AM, Urra X, van de Beek D, Veltkamp R. The immunology of acute stroke. Nat Rev Neurol. 2012;8:401–410.

23. Lin C-Y, Chang C, Cheung W-M, Lin M-H, Chen J-J, Hsu CY, Chen J-H, Lin T-N. Dynamic changes in vascular permeability, cerebral blood volume, vascular density, and size after transient focal cerebral ischemia in rats: evaluation with contrast-enhanced magnetic resonance imaging. J Cereb Blood Flow Metab. 2008;28:1491– 1501.

24. Fernández-López D, Faustino J, Klibanov AL, Derugin N, Blanchard E, Simon F, Leib SL, Vexler ZS. Microglial Cells Prevent Hemorrhage in Neonatal Focal Arterial Stroke. J Neurosci. 2016;36:2881–2893.

25. Bsat S, Halaoui A, Kobeissy F, Moussalem C, El Houshiemy MN, Kawtharani S, Omeis I. Acute ischemic stroke biomarkers: a new era with diagnostic promise? Acute Medicine & Surgery. 2021;8:e696.

26. Sladojevic N, Stamatovic SM, Johnson AM, Choi J, Hu A, Dithmer S, Blasig IE, Keep RF, Andjelkovic AV. Claudin-1-Dependent Destabilization of the Blood–Brain Barrier in Chronic Stroke. J. Neurosci. 2019;39:743–757.

27. Soontornmalai A, Vlaming MLH, Fritschy J-M. Differential, strain-specific cellular and subcellular distribution of multidrug transporters in murine choroid plexus and blood–brain barrier. Neuroscience. 2006;138:159–169.

28. Spudich A, Kilic E, Xing H, Kilic U, Rentsch KM, Wunderli-Allenspach H, Bassetti CL, Hermann DM. Inhibition of multidrug resistance transporter-1 facilitates neuroprotective therapies after focal cerebral ischemia. Nat Neurosci. 2006;9:487– 488.

29. Huang L, Chen Y, Liu R, Li B, Fei X, Li X, Liu G, Li Y, Xu B, Fang W. P-Glycoprotein Aggravates Blood Brain Barrier Dysfunction in Experimental Ischemic Stroke by Inhibiting Endothelial Autophagy. Aging Dis. 2022;13:1546–1561.

30. Kilic E, Spudich A, Kilic Ü, Rentsch KM, Vig R, Matter CM, Wunderli-Allenspach H, Fritschy JM, Bassetti CL, Hermann DM. ABCC1: a gateway for pharmacological compounds to the ischaemic brain. Brain: a Journal of Neurology. 2008;131:2679– 2689.

31. Manrique-Castano D, Sardari M, Silva de Carvalho T, Doeppner TR, Popa-Wagner A, Kleinschnitz C, Chan A, Hermann DM. Deactivation of ATP-Binding Cassette Transporters ABCB1 and ABCC1 Does Not Influence Post-ischemic Neurological Deficits, Secondary Neurodegeneration and Neurogenesis, but Induces Subtle Microglial Morphological Changes. Front. Cell. Neurosci. [Internet]. 2019 [cited 2025 Apr 1];13. Available from: https://www.frontiersin.org/journals/cellular-neuroscience/articles/10.3389/fncel.2019.00412/full

32. Dolman D, Drndarski S, Abbott NJ, Rattray M. Induction of aquaporin 1 but not aquaporin 4 messenger RNA in rat primary brain microvessel endothelial cells in culture. J Neurochem. 2005;93:825–833.

33. Kim JH, Lee YW, Park KA, Lee WT, Lee JE. Agmatine attenuates brain edema through reducing the expression of aquaporin-1 after cerebral ischemia. J Cereb Blood Flow Metab. 2010;30:943–949.

34. Roşu GC, Pirici I, Istrate-Ofiţeru AM, Iovan L, Tudorică V, Mogoantă L, Gîlceavă IC, Pirici D. Expression patterns of aquaporins 1 and 4 in stroke. Rom J Morphol Embryol. 2019;60:823–830.

35. Dorward HS, Du A, Bruhn MA, Wrin J, Pei JV, Evdokiou A, Price TJ, Yool AJ, Hardingham JE. Pharmacological blockade of aquaporin-1 water channel by AqB013 restricts migration and invasiveness of colon cancer cells and prevents endothelial tube formation in vitro. J Exp Clin Cancer Res. 2016;35:36.

36. Kanazawa M, Miura M, Toriyabe M, Koyama M, Hatakeyama M, Ishikawa M, Nakajima T, Onodera O, Takahashi T, Nishizawa M, et al. Microglia preconditioned by oxygen-glucose deprivation promote functional recovery in ischemic rats. Sci Rep. 2017;7:42582.

37. Kang M, Jin S, Cho H. MRI investigation of vascular remodeling for heterogeneous edema lesions in subacute ischemic stroke rat models: Correspondence between cerebral vessel structure and function. J Cereb Blood Flow Metab. 2021;41:3273– 3287.

38. Hu Y, Zheng Y, Wang T, Jiao L, Luo Y. VEGF, a Key Factor for Blood Brain Barrier Injury After Cerebral Ischemic Stroke. Aging Dis. 2022;13:647–654.

39. Moon S, Chang M-S, Koh S-H, Choi YK. Repair Mechanisms of the Neurovascular Unit after Ischemic Stroke with a Focus on VEGF. Int J Mol Sci. 2021;22:8543.

40. Hayashi T, Abe K, Suzuki H, Itoyama Y. Rapid induction of vascular endothelial growth factor gene expression after transient middle cerebral artery occlusion in rats. Stroke. 1997;28:2039–2044.

41. Li Z-R, Wang Y-Y, Wang Z-H, Qin Q-L, Huang C, Shi G-S, He H-Y, Deng Y-H, He X-Y, Zhao X-M. The positive role of transforming growth factor-β1 in ischemic stroke. Cellular Signalling. 2024;121:111301.

42. Taylor A, Verhagen J, Blaser K, Akdis M, Akdis CA. Mechanisms of immune suppression by interleukin-10 and transforming growth factor-beta: the role of T regulatory cells. Immunology. 2006;117:433–442.

43. Nishi T, Maier CM, Hayashi T, Saito A, Chan PH. Superoxide dismutase 1 overexpression reduces MCP-1 and MIP-1 alpha expression after transient focal cerebral ischemia. J Cereb Blood Flow Metab. 2005;25:1312–1324.

44. Zeng J, Wang Y, Luo Z, Chang L-C, Yoo JS, Yan H, Choi Y, Xie X, Deverman BE, Gradinaru V, et al. TRIM9-Mediated Resolution of Neuroinflammation Confers Neuroprotection upon Ischemic Stroke in Mice. Cell Reports. 2019;27:549–560.e6.

45. Sofroniew MV. Astrocyte barriers to neurotoxic inflammation. Nat Rev Neurosci. 2015;16:249–263.

46. Boveris A, Chance B. The mitochondrial generation of hydrogen peroxide. General properties and effect of hyperbaric oxygen. Biochem J. 1973;134:707–716.

47. Yamagata K, Tagami M, Takenaga F, Yamori Y, Itoh S. Hypoxia-induced changes in tight junction permeability of brain capillary endothelial cells are associated with IL-1beta and nitric oxide. Neurobiol Dis. 2004;17:491–499.

48. Sugawara T, Fujimura M, Noshita N, Kim GW, Saito A, Hayashi T, Narasimhan P, Maier CM, Chan PH. Neuronal death/survival signaling pathways in cerebral ischemia. NeuroRx. 2004;1:17–25.

49. Powanda DD, Chang TMS. Cross-linked polyhemoglobin-superoxide dismutase-catalase supplies oxygen without causing blood-brain barrier disruption or brain edema in a rat model of transient global brain ischemia-reperfusion. Artif Cells Blood Substit Immobil Biotechnol. 2002;30:23–37.

